# SIFNet: Electromagnetic Source Imaging Framework Using Deep Neural Networks

**DOI:** 10.1101/2020.05.11.089185

**Authors:** Rui Sun, Abbas Sohrabpour, Shuai Ye, Bin He

## Abstract

Electroencephalography (EEG) and magnetoencephalography (MEG) are used to measure brain activity, noninvasively, and are useful tools for brain research and clinical management of brain disorders. Tremendous effort has been made in solving the inverse source imaging problem from EEG/MEG measurements. This is a challenging ill-posed problem, since the number of measurements is much smaller than the number of possible sources in the brain. Various methods have been developed to estimate underlying brain sources from noninvasive EEG/MEG as this can offer insight about the underlying brain electrical activity with significantly improved spatial resolution. In this work, we propose a novel data-driven *Source Imaging Framework* using deep learning neural networks (SIFNet), where (1) a simulation pipeline is designed to model realistic brain activation and EEG/MEG signals to train generalizable neural networks, (2) and a residual convolutional neural network is trained using the simulated data, capable of estimating source distributions from EEG/MEG recordings. The performance of our proposed SIFNet approach is evaluated in a series of computer simulations, which indicates the excellent performance of SIFNet outperforming conventional weighted minimum norm algorithms that were tested in this work. The SIFNet is further tested by analyzing interictal EEG data recorded in a clinical setting from a focal epilepsy patient. The results of this clinical data analysis indicate accurate localization of epileptogenic activity as validated by the epileptogenic zone clinically determined in this patient. In sum, the proposed SIFNet approach promises to offer an alternative solution to the EEG/MEG inverse source imaging problem, shows promising signs of being robust against measurement noise, and is easy to implement, therefore, being translatable to clinical practice.

## 1 Introduction

### Conventional Approaches in Electrophysiological Source Imaging

Electroencephalography (EEG) and magnetoencephalography (MEG) are noninvasive techniques that measure synchronized activities of neuronal ensembles in the brain. Scalp EEG/MEG topography provides spatial information about the underlying brain electrical activity. However, such scalp topographical maps can only provide a rough estimation of the underlying sources due to volume conduction effect. Electrophysiological source imaging (ESI) was developed to significantly enhance the spatial resolution of noninvasive EEG/MEG. ESI is the process of estimating the underlying brain electrical activity from EEG (or MEG) recordings, through a deconvolution process, to find the brain source distributions that fit the recorded EEG (or MEG) [1]. However, it is challenging to localize and image the brain electrical activity precisely and reliably with only a few hundreds of sensors, because of the ill-posed nature of the inverse problem; as different source distributions can produce the same potentials/fields at the scalp [1,2].

To overcome the challenges posed by the limited number of measurements and the volume conductor effect, current ESI methods require a priori assumptions, albeit sometimes reasonable, to regularize the inverse problem and to ultimately find a unique solution [3]. The dipole source localization methods need to know the number of sources a priori [4], and when that information is not available, distributed-source models can be used [5] with regularization priors to restrict the solution space. Regularization terms could be based on imposing constraints on solutions’ energy [6–8], covariance [9,10], sparsity level [11–13] or multiple different constrains [14]. These methods require parameter tuning to balance between fitting the recorded data and satisfying the regularization constraints when solving every instance of a given problem, i.e. for every given EEG/MEG measurement [15], which could be a time-consuming process.

### ESI Approaches based on Deep Learning

Deep learning approaches aim at capturing the correct mapping between signal and source spaces through a large amount of training data which is ultimately represented by the weights and non-linear units in its layers. Another feature of deep learning approaches is that after the time-consuming training process, the algorithm is quite efficient and fast when applied to new data. It spares the need to search for the hyperparameters for each new instance of EEG/MEG measurement, therefore providing an opportunity for real-time source imaging.

There have been a few attempts to solve the dipole fitting problem for one or two dipoles using neural network approaches decades ago [16–20]. These works reported a wide range of performance levels in simulation from a localization error of 3 cm [16] to 3.5 mm [17]. Abeyratne et al [20] tested the trained neural network on the data recorded from epilepsy patients, and showed the normalized mean square error of reconstructed EEG signal was around 20%. However, those solutions were obtained with the a priori knowledge of number of sources, and brain sources were modeled as one or two current dipoles.

### Realistic Modeling of Dynamic Brain Activities

To develop a successful analysis tool using deep learning, the neural network algorithms need to be trained successfully. For model training, it is crucial to obtain enough labeled data to provide a proper mapping pattern from input to output. Since the amount of EEG/MEG data with source location correctly labeled is usually limited [21], computational models can be used to generate the training data, as well as the testing data, to quantitatively evaluate the model performance.

Many computational methods, from microscopic to macroscopic level, have been proposed to model the various kinds of neural activity, e.g. alpha oscillations vs. inter-ictal spikes, and many can be adopted to generate the training data for neural networks. One approach is to model the brain electrical activity using equivalent physical models, such as few current dipoles [4] or distributed current density models [6]. In such equivalent physical source models, other regions are treated as independent units, and are assigned with different temporal activity profiles to distinguish their electrophysiological functions. For instance, the cortical regions which is the source of inter-ictal activity will have a temporal profile that resembles a spike and other regions will have Gaussian noise-like activity. These physiologically plausible models are widely used in ESI studies [22,23]. Prior efforts using neural network based approaches also used such equivalent physical source models (one or two dipoles) to generate their training dataset [16–20].

On the other hand, there are models that take the neuronal dynamics based on cellular modeling and the interaction between different brain regions into account, and the source activities of the whole brain are generated from these interactions. One well-studied model is the neural mass model (NMM) which is based on neurophysiological principles [24–26]. NMM models the average electrical activity of principal neurons and interneurons ensembles [27] of a cortical region, and not individual neuron’s activity, hence operating on a mesoscopic scale. The temporal behavior of the model is defined by differential equations and model parameters, such as tissue properties, e.g. excitability, and inter-regional connectivity [28]. These models run on noise and once a few parameters are set, they can autonomously generate EEG-like (or MEG-like) activity. It was successfully applied in generating rhythmic activity [29], event-related potentials [30], and was further modified to model interictal spikes [31]. NMM has been used in conjunction with the forward modeling of electromagnetic field propagation in the head tissue, i.e. EEG and MEG lead-field, to provide more insight into the underlying pathophysiological mechanisms of epileptic activity [27,32].

### ESI by Means of Deep Learning and Realistic Brain Modeling

In the proposed SIFNet approach, we integrate the deep learning neural network with the realistic dynamic brain models such as NMM, to provide a framework that the forward source-signal relationship can be modeled using neural mass models, and inverse source imaging can be performed by means of deep learning.

In this work, we develop a *S*ource *I*maging *F*ramework that uses dynamically generated data by means of NMM models to train a convolutional neural *Net*work for distributed source imaging, which we call for short SIFNet. This is the first report, to our knowledge, to use realistic neural mass model generated data to train an artificial neural network solving the source imaging problem capable of providing distributed solutions. We illustrate the merits and applicability of the proposed SIFNet approach to solve epilepsy source localization, which serves as our testing benchmark without attempting to model all possible brain dynamics in its entirety, given the challenges involved. We show: (1) when the neural network is trained with realistic simulation brain activities generated by interconnected NMM networks, it can outperform conventional ESI methods; (2) SIFNet’s superior performance can be generalized to signals simulated under different protocols; and (3) its potential on solving the ESI problem from real EEG data, as showcased by analyzing epileptic interictal spikes.

## 2 Methods

While the proposed SIFNet approach can be applied to both EEG and MEG data, for the simplicity, we will only show its application to EEG data in the following. As illustrated in Figure 2, NMM generated signals are used as the underlying brain activities and the simulated scalp EEG data was obtained by multiplying the generated NMM signals by the lead-field matrix. The synthetic scalp EEG measurements were used as input to SIFNet during the training process. The network was updated iteratively to learn the mapping relationship between the sensor space and the source space. Lastly, three sets of data with different source signal distributions were tested on the trained network.

**Figure 1.**
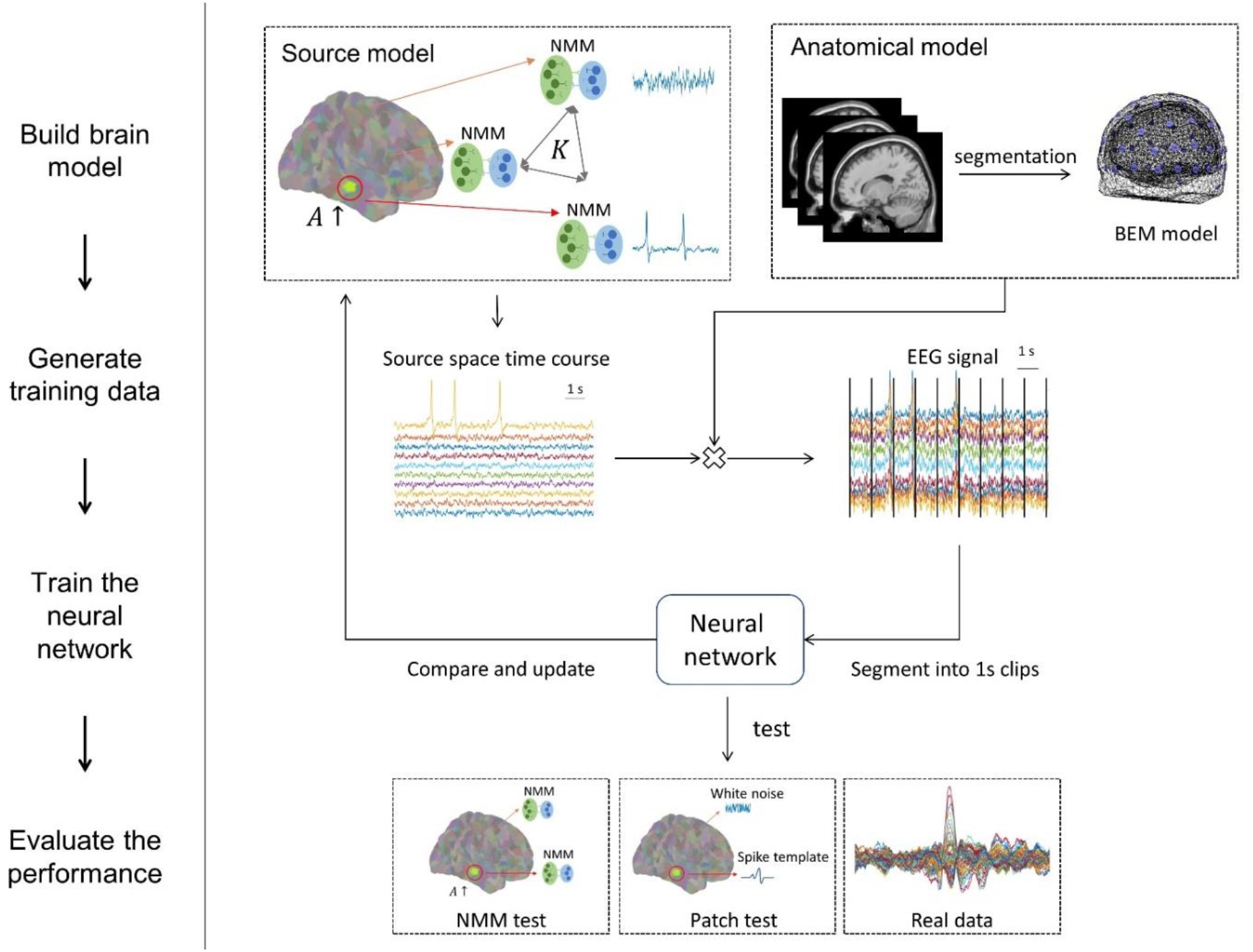
Source imaging framework using neural networks. The structure of this study and the testing scenarios used here is to address the 3 fundamental questions raised, namely, how to generate enough training data that resembles real EEG, how robust a model trained on these models will be compared to variations in the input data, and how well will a model trained on these premises perform in real-EEG recordings.

**Figure 2.**
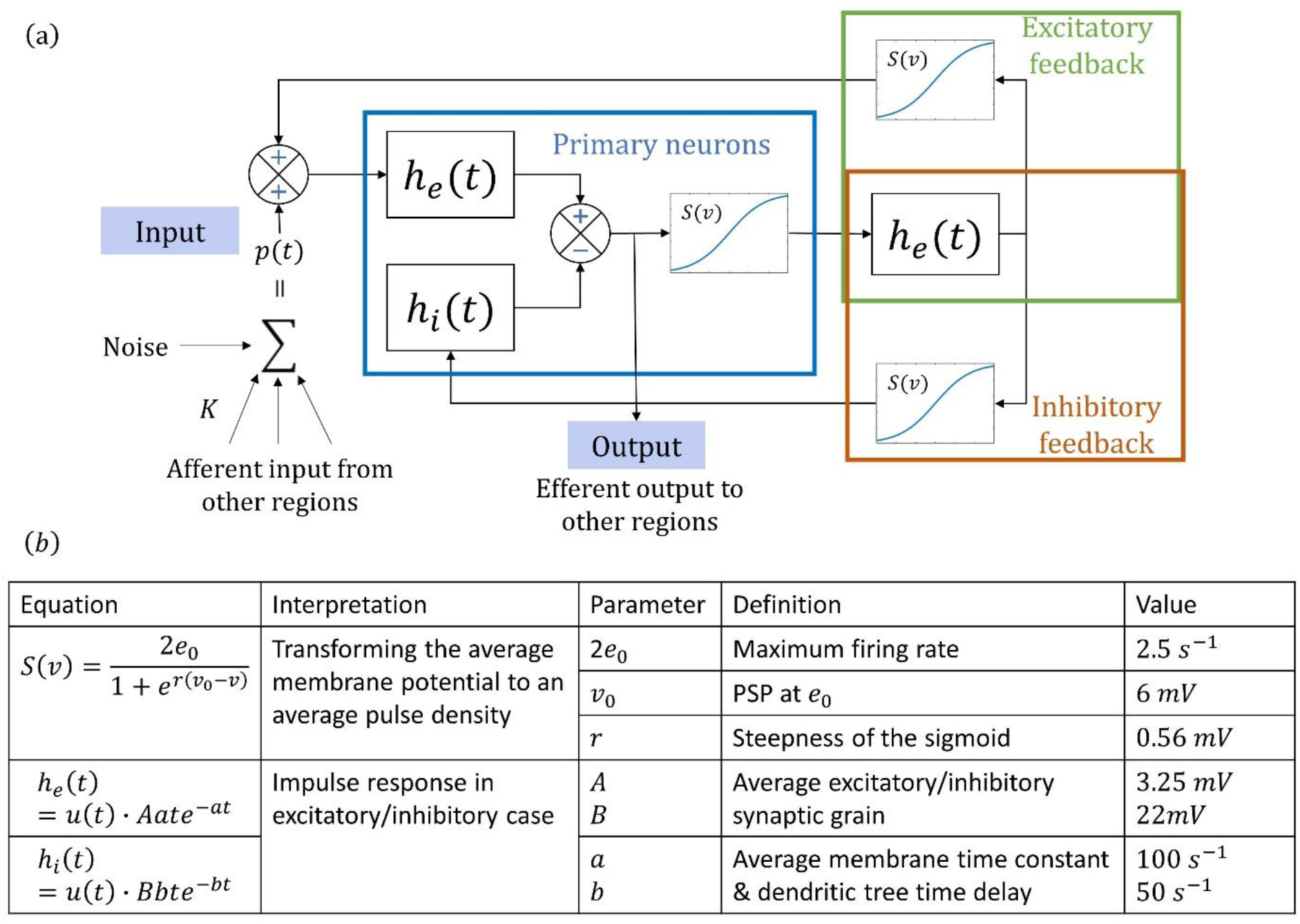
(a) Structure of a single-element Jansen-Rit model. A single-element Jansen-Rit model simulates three neuron sub-populations: the primary neurons (i.e. pyramidal cells), as well as excitatory and inhibitory interneurons. The influence from other neural populations is modeled as an excitatory input p(t) that represents the average pulse density, i.e. average firing rate, of afferent action potentials. The behavior of each sub-population is modeled using linear functions h_e (t) and h_i (t) that describe how presynaptic information is transformed into postsynaptic information, and nonlinear function S(v) that models the change from average membrane potential of the subpopulation to average pulse density. (b) Equations and parameters defined in a single-element Jansen-Rit model and their interpretation. Adapted from [31].

### 2.1 Modeling Temporal Dynamics with Networks of Neural Mass Models

The temporal dynamic of the brain activity can be generated using interconnected neural populations, each represented by a physiologically relevant model. The Jansen-Rit model [30] was shown to be a reasonable model for interictal spike activities [33,34]. The source activity of each brain region is simulated by modeling the interaction of neuron sub-populations through differential equations. The relationship between neuron sub-populations are illustrated in Figure 2(a). The equations and the standard values for the parameters are listed in Figure 2(b).

With the standard values provided in Figure 2(b), the NMM network can simulate brain signals similar to resting-state brain activity, especially in the alpha band [30], and when the average excitatory synaptic gain (defined as *A*) is increased, it can simulate spike-like activities, which exhibit similarities with interictal spike signals observed in real recordings [31]. Many single-element Jansen-Rit models can be connected through a connectivity matrix *K*, where the input for each single element is the weighted sum of other elements’ output to form a network of interconnected NMMs. By increasing *A* for one NMM and keeping the other *A* values the same, we can simulate the temporal dynamic of the brain with one “hyperexcitable” region, resembling the simplified dynamics observed in focal epilepsy patients [33].

### 2.2 EEG Signal Simulation

A template T1-weighted MRI was used to get the segmentation of the cortical surface, skull and skin using Freesurfer [35]. The 64-channel BioSemi electrode layout (BioSemi, Amsterdam) was used for the EEG electrode configuration. The forward matrix was calculated with the conductivity values of 0.33, 0.0165 and 0.33 S/m for the inner skull, outer skull and scalp, respectively [36], using the boundary element method (BEM) model [4,37]. The source space was defined over the cortex by segmenting the whole cortex into 994 similarly sized interconnected regions with a current dipole source located at the center of each region whose time course was generated by the NMM. The connectivity weights *K* between those regions were calculated from diffusion-weighted images averaged over 5 healthy subject as provided in [38]. The NMM network modeling was conducted in *The Virtual Brain* simulator with a sampling rate of 500 Hz [32].

The brain source signals from all regions were then projected to the scalp using the lead-field matrix. The NMM signal from each region was scaled so that the ratio between the interictal spike signal, and the background signal from other regions contributing to the EEG, was the same for all simulated data. In other words, cortical regions which had inherent disadvantage due to their depth were normalized so that they received equal representation in the simulated EEG as regions which had stronger representation. Afterwards, different levels of Gaussian white noise were added to the scalp potential to simulate noise-contaminated EEG measurements, so that the SNR between the EEG signal and the white noise was 5, 10, 15 or 20 dB, respectively. The signal was then segmented into 1-second clips and fed as the input into the neural network. In total, the training data contained about 500,000 extracted spike intervals with different source distributions and noise levels. The protocol for the data-generation process is shown in Figure 1.

The test data were separately generated following the same protocol. However, the mean and variance for the noise input in p(t) (see Figure 2) were slightly different from the training data. Since the noise input influences the state of the differential equations that governs NMM outputs, changing the noise will alter the signal dynamic. By applying these perturbations the test data were not identical to the training data. About 40,000 testing examples were simulated. The training and testing data generated using the NMM network are called *nmm train* and *nmm test*, respectively, from now on.

#### An Alternative Method for Source Simulation

In addition to NMM-generated source signals, an alternative method was employed to simulate these sources by using pre-defined signal templates. Since this study focused on epileptic interictal spike localization applications, we used four different interictal spike signals extracted from EEG recordings of epilepsy patients. Then we assigned these extracted signals to the source region serving as the spike generating cortical region and added Gaussian white noise to the other regions so as to simulate the effect of the colored noise observed in EEG from other brain regions, similar to NMM-generated data. After the brain signal was projected onto the scalp, different levels of Gaussian white noise were also added to the simulated EEG. For the testing data in this scenario, a different spike template was used. The same amount of training and testing data was generated, as the NMM simulation protocol. The data generated in this scenario are named *patch train* and *patch test*, respectively.

### 2.3 SIFNet Architecture

The spatiotemporal EEG data are represented as a matrix of channel cross time. Convolutional neural network (CNN) was applied to process the input. CNN has shown superior performance in many EEG-related deep learning applications [39]. It can extract local, common features by using convolution and nonlinear activation functions, and global features can be learned hierarchically by layered computations [40]. Since the activity arising from each source location contributes to all EEG channels, the source activity can be fully estimated if global features are considered and analyzed; a capability that CNN architecture provides. To achieve the goal of ESI, which is to estimate underlying brain activities at the source-level, it is important that there are enough convolution filters that have wide-enough receptive fields covering all EEG channels [41]. In other words, to aggregate the local spatiotemporal features and unmix the global source information, a deep enough neural network is needed.

To mediate the difficulties in training deep neural networks, a residual structure was adopted [42]. Five residual blocks were followed by a pooling layer and a fully connected layer with its output size the same as the number of source regions. Each residual block contained two CNN layers with batch normalization to remove the covariance shift during the training [43]. The original input was added back to the output and fed to the next block through a Rectified Linear Unit (ReLU) activation function [44]. The ESI problem was reformulated as a classification problem and cross entropy between the true source region, i.e. spike generating region, and the predicted source region was used as the loss function. Adam optimizer [45] was used for the training. The whole network was implemented in PyTorch^1^. The detailed network structure is illustrated in Figure 3.

**Figure 3.**
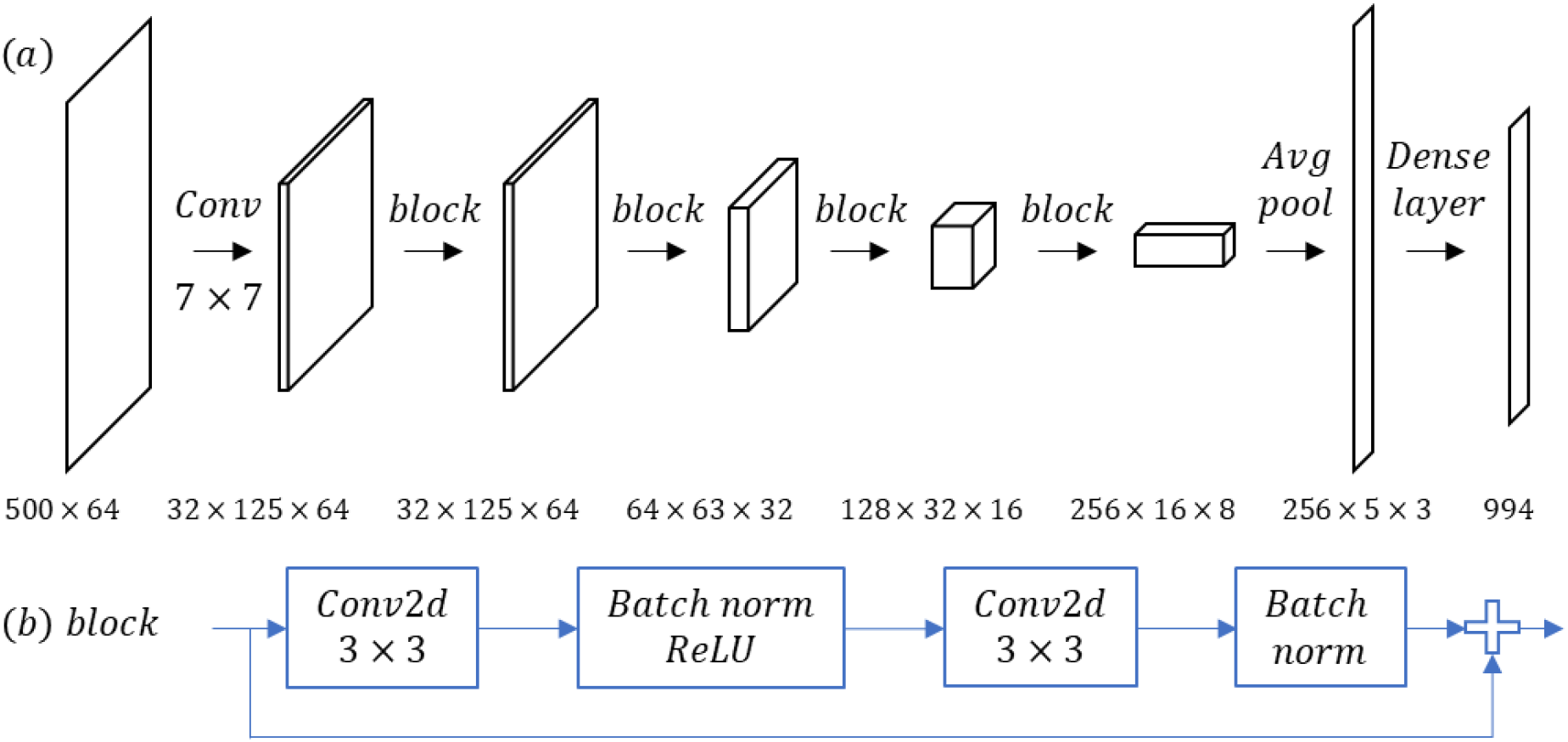
The diagram for the SIFNet network architecture. Conv2d means 2-dimensional convolution and *n* × *n* indicates the convolution kernel size. It has about 5 million parameters in total. (a) The whole network structures. (b) The computation inside the residual block.

### 2.4 Evaluation Protocol

Two models with the same architecture as described above were trained with either *NMM train* or *patch train* data till convergence. The acquired models were named as *NMM trained model* and *patch trained model*, respectively. Both models were tested on *NMM test* and *patch test* to evaluate its generalizability on different source dynamics. The results on both test datasets were further compared to one popular conventional ESI method: standardized low resolution electromagnetic tomography (sLORETA) [6] (we employed the Brainstorm [46] implementation of the algorithm). In a word, three different methods, *NMM trained model, patch trained model* and sLORETA, were tested on two different datasets *NMM test* and *patch test*.

The performance was evaluated using two different metrics, namely, the localization error and the concordance rate. The localization error (LE) was defined as the Euclidian distance between centers of the true source and the predicted source. Concordance rate was defined as the percentage of cases of the spiking region identified by the source localization result that overlapped with the true simulated source. Wilcoxon rank sum test implemented in MATLAB was used to investigate statistical significance of the obtained results.

## 3 Evaluation Results

The two columns in Figure 4 (a) represent the overall LE for all noise levels on *patch test* and *NMM test* for three methods: *NMM trained model*, *patch trained model*, and sLORETA. Each model was trained with 500,000 samples and each test dataset includes a total of 40,000 examples, covering four different noise levels. The solid line shows the mean and standard deviation, the dotted line shows the median and the color bar spans the 90th percentile of the data. The distribution of localization errors is shown in Figure 4(b).

**Figure 4.**
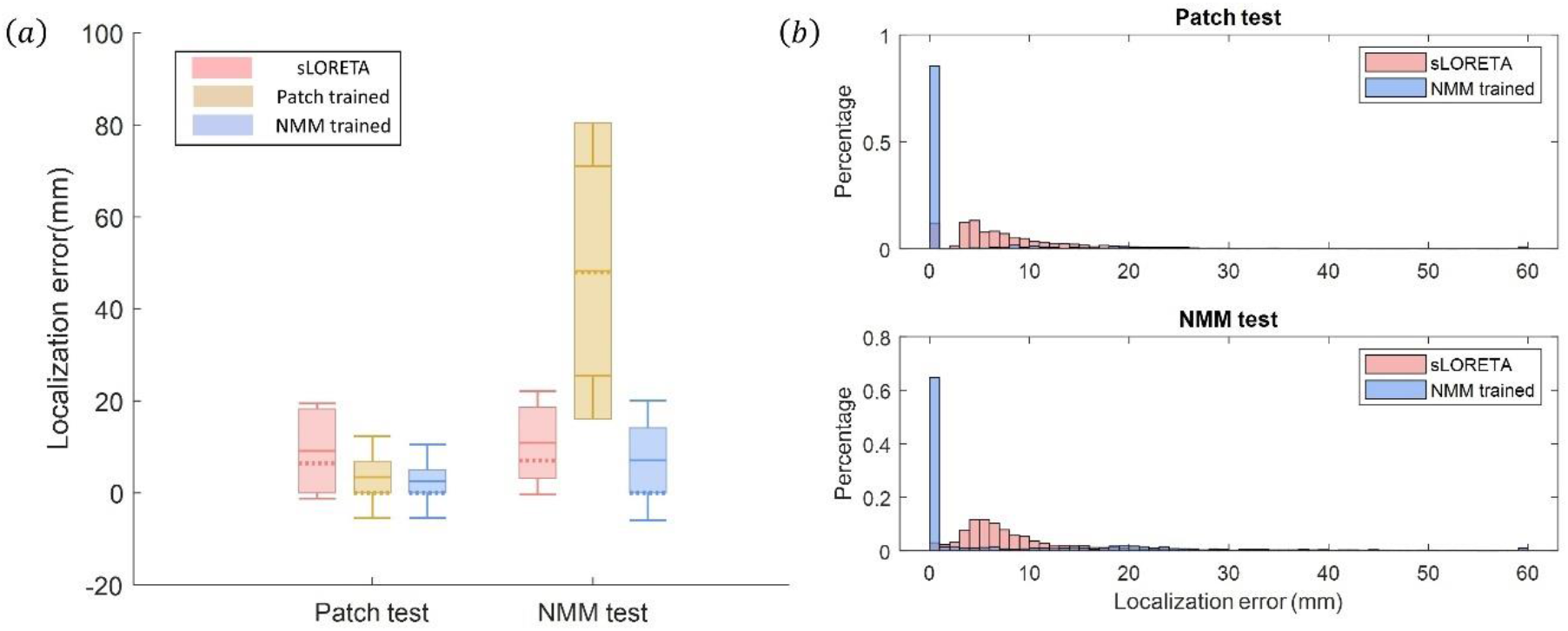
(a) Localization error of NMM trained model, patch trained model and sLORETA for *patch test* and *NMM test*. Each model was trained with 500,000 samples and each test dataset includes a total of 40,000 examples. The localization error (LE) was defined as the Euclidian distance between the true source and the predicted source. The solid line shows the mean and standard deviation, the dotted line shows the median and the color bar spans the 90% percentile of the data. (b)The localization error histograms of sLORETA and NMM trained models. For better visualization, LE above 60 mm is projected to the 60 mm bin.

### *NMM Trained Model* vs *Patch Trained Model*

In *patch test*, where the data was generated using Gaussian white noise and independent dipoles, we can see *NMM trained model* and *patch trained model* reached similar mean localization error (Mean ± standard deviation: *Patch trained model:* 3.43 ± 8.91 mm; *NMM trained model:* 2.53 ± 7.97 mm; p < 0.01). It is expected for the *patch trained model* to perform well on *patch test*, since the training and testing data were generated using the same protocol. However, we can see that even though the NMM trained model is trained on a different dataset, it demonstrates robustness and generalizability when tested on a different dataset.

In *NMM test*, where the data were generated by interconnected NMMs, the *NMM trained model* outperformed the *patch trained model* significantly (*NMM trained model* 7.07 ± 13.06 mm; Patch trained model: 48.24 ± 22.74 mm, p < 0.01). As we can see from Figure 4(a), the *patch trained model* barely produced meaningful results in the test, which meant it would not work on more elaborate scenarios where background brain activity or noise are correlated, which is highly likely in reality.

### *NMM Trained Model* vs sLORETA

We also compared the NMM trained model with a conventional source imaging method, sLORETA. In terms of the localization error, as we can see in Figure 4(a), *NMM trained model* has a lower mean value (*patch test: NMM trained model* 2.53 ± 7.97 mm; sLORETA 9.12 ± 10.33 mm; p < 0.01; nmm test: *NMM trained model* 7.07 ± 13.06 mm; sLORETA 10.89 ± 11.24 mm; p < 0.01). The *NMM trained model* also showed a very high concordance rate of 87% compared to the 53% of sLORETA. This can be observed from the histogram of localization errors in Figure 4(b), where for *NMM trained model*, most of the test data have a localization error of 0 for both test datasets as the model identified the correct region, while sLORETA has distributed error values between 0 to 20 mm.

In order to explore what most impacts the model performance, the performance of the two approaches under different SNR and depth conditions (how deep the active source is located from scalp) is calculated and summarized in Figure 5. The SNR values represent the level of Gaussian white noise added to the EEG signal when generating the data. The depth was defined as the minimum distance between the source and the closest scalp electrode. Localizing sources with noisy EEG recordings, or deeply located sources has always been a challenging task in the ESI field [47–49], as evident in Figure 5; the LE increases with the increasing depth of underlying sources and decreases as SNR increases, for sLORETA estimates. The LE for NMM trained models, however, is flat under these conditions. This we believe is because of the fact that the deep learning model had learned the mapping rules under deep-source and low-SNR conditions, through the training process, so its performance was not negatively impacted by these challenging conditions.

**Figure 5.**
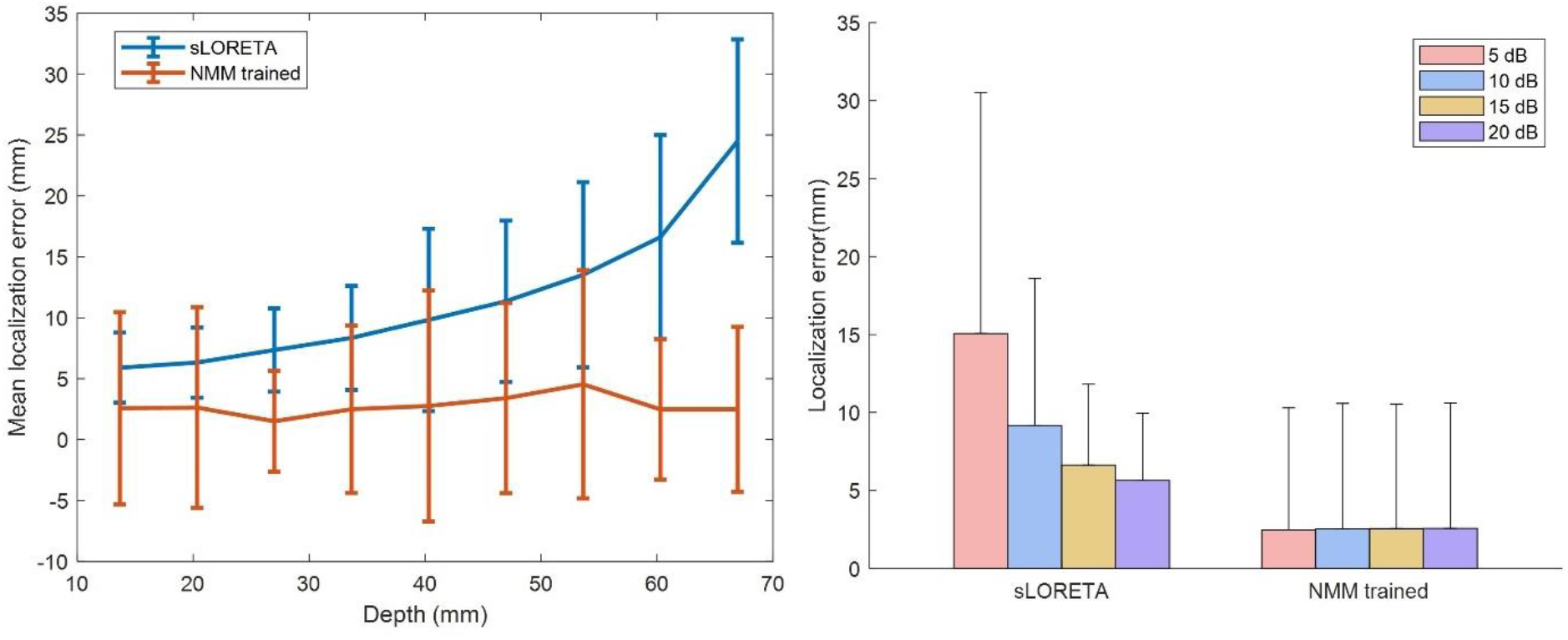
The influence of depth and noise on localization error. Left: The mean localization error vs. source depth. Depth is defined as the minimum distance between the scalp and the source location. Error bar shows the standard deviation. The difference between sLORETA and NMM for different SNR levels. Bar shows the mean of LE at different SNR level and line shows the standard deviation.

## 4 Real Data Analysis

All the methods, *NMM trained model, patch trained model* and sLORETA were also tested on the recordings from one patient that suffered from right temporal lobe epilepsy, who became seizure-free after surgical removal of the epileptogenic tissue. The patient’s scalp EEG recordings was resampled to 64 channels (same size as the SIFNet inputs), and was bandpass filtered between 0.5 and 40 Hz. 58 interictal spikes were identified and analyzed in the EEG recordings of this patient. An example spike signal and its topography at the peak are shown in Figure 6. Patient’s MRI was co-registered with the template MRI (used to generate the training data). ESI was performed on each individual spike waveforms, and the average result was used to evaluate the performance.

**Figure 6.**
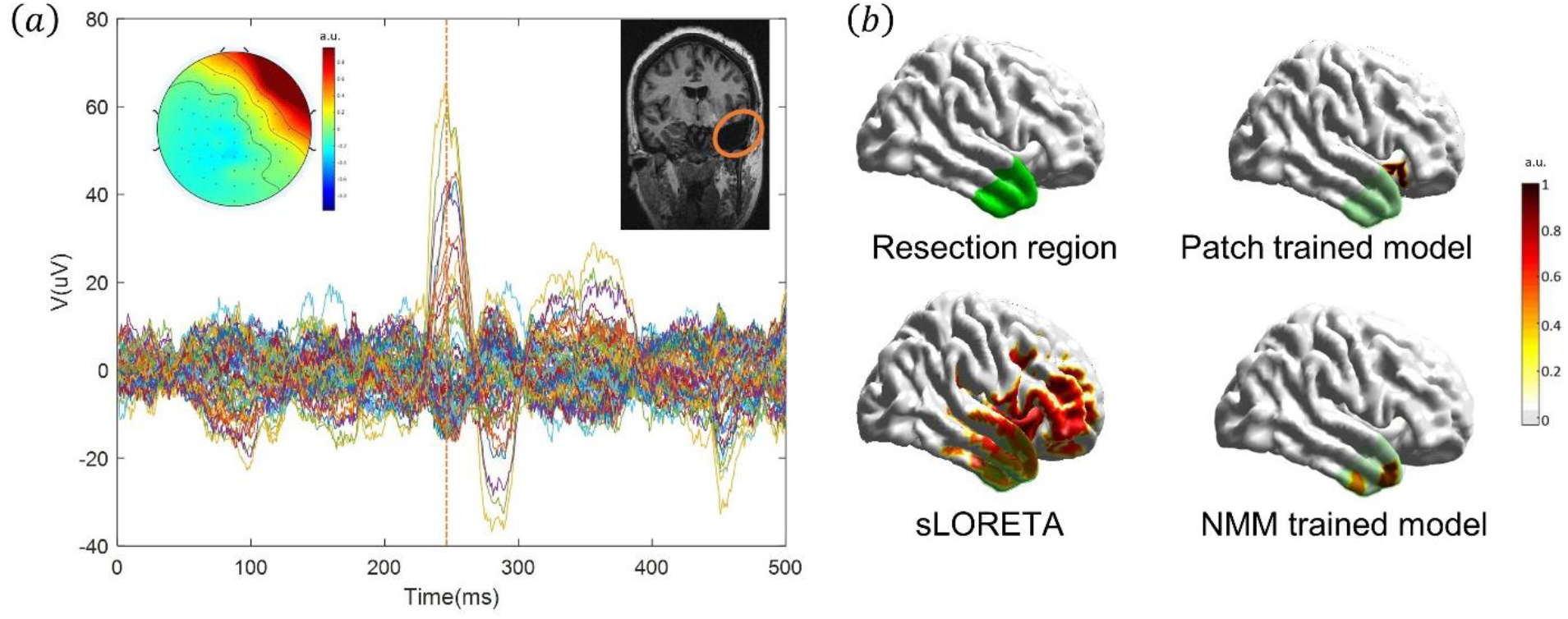
Epilepsy patient data analysis. (a) Butterfly plot of the EEG waveform of an example spike analyzed, with topography on the peak of the spike, and the illustration of surgical resected area. (b) Comparison of source imaging results using NMM trained model, patch trained model, and sLORETA. The green color indicates the resection area and the red color is the identified source region by the models. For patch trained model and NMM trained model, the color-bar shows the normalized number of spikes localized into a certain region, and for sLORETA, the color-bar shows the normalized scale of solution amplitude of the averaged source reconstructions.

The test results on the interictal spike data from the epilepsy patient are shown in Figure 6. The *patch trained model’s* result was on the inferior frontal gyrus, outside the epileptogenic zone. The maximum of sLORETA’s result was located inside the resection region, however, the sLORETA estimate had some spurious activities on the frontal lobe. The source region identified by the *NMM trained model* was at the anterior portion of the temporal lobe, which lied inside the resection area as shown in Figure 6(b). Fig. 6 suggests that SIFNet can localize epileptiform activity concordant with the clinically identified source regions without having nuisance spurious activities.

## 5 Discussion

ESI has been shown to significantly improve the spatial resolution of scalp recorded EEG, yet, due to the challenges posed by its ill-posed nature, more development in this field is necessary. Instead of adding explicit constrains and assumptions into the inverse methods, data-driven approaches like deep learning can learn the mapping relationship between the EEG signals and the source activities through large amounts of training data. Given the lack of large amount of EEG recordings with corresponding source information, synthetic data are needed for deep learning based ESI studies.

In this work, we have proposed the SIFNet, a novel data-driven source imaging framework where realistic brain and EEG signals are simulated for training neural networks, and a residual convolutional neural network is trained using the synthetic data. In simulation evaluation and real patient testing, the proposed framework demonstrated excellent and consistent results under different signal conditions and proved to be robust when tested and trained on different models.

### The Importance of Generating Realistic Simulation Data

Ensuring generalization of deep learning models is a challenging and on-going research area [50]. It becomes more challenging when the training data and the testing data come from different data distributions. The performance of any deep learning based ESI method depends heavily on the similarity between its training data and the real data it ultimately has to analyze. In other words, the extent to which simulated training data represents the practical testing data, influences the performance of deep learning based ESI approaches as the model has to learn the characteristics of data it has to analyze through its training data.

This work shows the importance of having a realistic synthetic training dataset by testing the robustness of the same network architecture on two different sets of simulation data. As shown in Figure 4, *patch trained model* failed on *NMM test*, which means it cannot properly model the variabilities in other brain regions. This may introduce error when tested on real data, where the assumption that the brain signal in each region is independent Gaussian white noise, does not hold. On the other hand, the fact that *NMM trained model* has consistent results over the two tested datasets, shows that the commonly used white noise model to simulate background brain activity, i.e.*patch test*, was not enough to cover different test situations for data-driven methods, and it was necessary to use more realistic data generation methods, such as NMM networks, to train the neural networks for the purposes of ESI.

It is worth noting that the *NMM trained model* did not overfit onto NMM generated signals, but actually learned the mapping relationship between brain sources and EEG signals. *NMM test* itself is a more complex dataset with signals generated by interconnected brain regions, so it is expected that the performance is worse compared to *patch test*. If the *NMM trained model* was overfitted to the train data, it would have a much lower LE. The fact that it had larger LE on *NMM test* compared to *patch test*, aligns with the expectation and provides another evidence of its generalizability.

### SIFNet Can Handle More Challenging Situations

Conventional model-driven methods perform ESI based on the information from signal test instance and the priors defined in the algorithms. Their performance tends to suffer when the signal quality is low, when sources are located in deep regions or in low-SNR conditions [49]. With data-driven approaches, the correct localization under these conditions is provided to the neural network during the training phase, and the mapping function is implicitly stored in the weights in the neural network. As a result, SIFNet has stable and consistent performance across different source depth and SNR levels as shown in Figure 5.

### Limitations

In the current framework, only one-source configurations were tested. That is at a given time, only one spiking region existed in the simulation scenarios. However, SIFNet can extend to include multiple-source configurations by including multiple simultaneously active sources in the training data.

Estimating the underlying sources’ extent is also an important issue that needs further investigation. Some recent developments in the field suggest that underlying source sizes can be reliably estimated from noninvasive scalp measurements such as EEG [13]. Additionally, estimating the temporal dynamics of estimated sources, rather than just identifying the location of activity in the brain, is of utmost importance when studying a dynamic system like the brain. These objectives call for more accurate source models, deep learning models with more layers, and more elaborate loss functions.

On a more practical note, to apply SIFNet to analyze a large cohort of patients’ data, the inter-subject variabilities could have an impact on the model performance; these include conductivity uncertainties, sensor configuration mismatches, anatomical variabilities, etc. These factors impair the performance of conventional methods, as well, and their effects on SIFNet need further careful examination [36,51]. There are multiple possible ways to incorporating these variabilities into the framework. Acquiring individual neural networks using the subjects’ own anatomical information is straightforward but time-consuming option. In this approach one trains the SIFNet for each individual subject based on the subject’s individual head model. One could also fine-tune the model by continuously training the model with data from each individual in the data base, with the expectation that the final trained model captures individual anatomical variabilities becoming robust to such perturbations.

In sum, further investigations are needed and work can be extended to incorporate various practical factors into the current framework to provide a stable and robust ESI method translatable to real-world and clinical settings.

## ACKNOWLEDGEMENTS

This work was supported in part by NIH grants EB021027, NS096761, EB029354, MH114233, AT009263.

This work used the Extreme Science and Engineering Discovery Environment (XSEDE), which is supported by National Science Foundation grant number ACI-1548562. Specifically, it used the Bridges system, which is supported by NSF award number ACI-1445606, at the Pittsburgh Supercomputing Center (PSC).

## Footnotes

1 https://pvtorch.org/

